# Substitution of a valine to glutamic acid in the omega-like loop of MSMEG_6194 of *Mycobacterium smegmatis* interchanges its activity from DD-carboxypeptidase to beta-lactamase

**DOI:** 10.1101/2025.11.20.689485

**Authors:** Aditya Prasad Panda, Debasmita Chatterjee, Anik Roy, Sarmistha Biswal, Anindya S Ghosh

**Affiliations:** Department of Bioscience and Biotechnology, Indian Institute of Technology Kharagpur, West Bengal, India, PIN-721302; Department of Infectious Disease Biology, Institute of Life Sciences, NALCO Square, Chandrasekharpur, Bhubaneswar, Odisha, 751023, India

**Keywords:** MSMEG_6194, Mycobacteria, class A beta-lactamases, bocillin-FL, penicillin-interacting protein

## Abstract

The genome of *Mycobacterium smegmatis* encodes numerous penicillin-interacting enzymes, and analysing their functions provides insights into the evolutionary mechanisms behind beta-lactam resistance in mycobacteria. In this study, we characterised one such enzyme, MSMEG_6194, annotated as a putative beta-lactamase. Although MSMEG_6194 shares structural similarity with class A beta-lactamases, it showed no detectable beta-lactamase activity under the tested conditions. Heterologous expression of MSMEG_6194 in *Escherichia coli* and *Δmsmeg_6194* deleted strains of *M. smegmatis* did not confer significant resistance to beta-lactams, and the purified protein failed to hydrolyse nitrocefin either. However, ectopic expression of MSMEG_6194 partly restores the morphological defects in seven PBP-deleted *E. coli* strains, and the purified enzyme successfully cleaves the terminal D-alanine from a pentapeptide substrate, confirming its DD-carboxypeptidase activity. Structural analysis revealed the absence of a conserved glutamic acid residue in the omega-loop, which is critical for beta-lactamase catalysis in class A beta-lactamase. Substituting this residue (V139E mutant) imparts beta-lactamase activity though significantly reduces DD-carboxypeptidase function. Overall, these findings establish MSMEG_6194 as a DD-carboxypeptidase and demonstrate how a single amino acid change can alter catalytic preference, shedding light on the evolutionary transition from DD-Carboxypeptidases to beta-lactamases in mycobacteria.

## Introduction

The bacterial cell wall is crucial for maintaining cellular shape and integrity, with peptidoglycan serving as its major structural component. The synthesis, maintenance, and regulation of peptidoglycan are orchestrated by a set of enzymes collectively known as penicillin-binding proteins (PBPs) (Ghosh et al. 2008, Sauvage et al. 2008). These enzymes are broadly classified into high-molecular-mass (HMM) and low-molecular-mass (LMM) PBPs, which perform diverse catalytic functions, including transpeptidase, transglycosylase, and carboxypeptidase activities. While LMM-PBPs predominantly exhibit DD-carboxypeptidase activity, some members also display transpeptidase or endopeptidase functions (Ghosh et al. 2008).

The beta-lactam antibiotics target PBPs by covalently acylating the active-site serine residue, thereby irreversibly inactivating the enzyme (Zapun et al. 2008). However, bacteria have evolved mechanisms to resist these antibiotics, either by modifying their PBPs or by producing beta-lactamases (BLs) that hydrolyse and neutralise the drug (Mora-Ochomogo & Lohans 2021). Notably, PBPs and beta-lactamases share a common evolutionary origin, giving rise to multiple classes of enzymes with distinct yet related catalytic mechanisms (Massova & Mobashery 1998). Based on sequence homology, beta-lactamases are grouped into four molecular classes (A–D). Classes A, C, and D are serine beta-lactamases, whereas class B enzymes are metallo-beta-lactamases (Bush & Jacoby 2010). Among these, class A beta-lactamases are evolutionarily closest to LMM-PBPs and share the conserved motifs SxxK, SxN, and KTG, which define their catalytic core (Massova & Mobashery 1998). Both enzyme types form an acyl–enzyme complex upon beta-lactam binding though, they differ in their deacylation efficiency. In PBPs, the acyl–enzyme intermediate remains stable, whereas in class A beta-lactamases, a glutamic acid residue within the omega-loop activates a water molecule, promoting deacylation and restoring enzyme activity, thereby completing the hydrolysis of antibiotics (He et al. 2020, Pemberton et al. 2020).

Despite their mechanistic similarities, the evolutionary divergence between PBPs and beta-lactamases remains incompletely understood. Most beta-lactamase studies have focused on enzymes from clinical isolates, leaving the natural diversity of these enzymes largely unexplored (Bush 2018). Investigating penicillin-interacting enzymes (PIEs) from non-pathogenic organisms can therefore provide us with valuable insights into the evolution of beta-lactam resistance mechanisms.

In this study, we functionally characterised the putative beta-lactamase gene *msmeg_6194* from *Mycobacterium smegmatis*. Although annotated as a beta-lactamase-related protein and structurally similar to class A beta-lactamases, our biochemical and physiological analyses demonstrate that MSMEG_6194 functions predominantly as a DD-carboxypeptidase in both *in vivo* and *in vitro*, and lacks detectable beta-lactamase activity, which is likely due to the absence of a critical glutamic acid residue within the omega-loop.

## Materials and Methods

### Bacterial strains, plasmids, and reagents

The bacterial strains and plasmids used in this study are outlined in Table S1 (supplemental material). *Mycobacterium smegmatis* mc^2^155 was cultured in Middlebrook 7H9 broth and 7H11 agar medium (Sigma-Aldrich, MO, USA), which was supplemented with 10% OADC, 0.35% (w/v) glycerol, and 0.05% (w/v) Tween 80. *Escherichia coli* strains were grown in LB medium. MH (cation-adjusted) media was used for the antibiotic susceptibility assay in *E. coli*. Antibiotics are used at the following concentrations: hygromycin at 100 µg/mL (Invitrogen, CA), kanamycin at 50 µg/mL, chloramphenicol at 20 µg/mL, and tetracycline at 0.02 µg/mL (HiMedia, Mumbai, India). All enzymes for genetic modifications were purchased from New England BioLabs (MA), and all reagents and chemicals were obtained from Sigma-Aldrich (St. Louis, MO) unless otherwise indicated.

### Recombinant plasmid construction and site-directed mutagenesis

The open reading frame (ORF) of *msmeg_6194* was amplified from the genomic DNA of *Mycobacterium smegmatis* mc²155 and cloned into the pBAD-18Cm vector using *NheI* and *HindIII* restriction sites to generate the construct pD6194. For expression in *M. smegmatis*, the same ORF was cloned into the pMIND vector using *NdeI* and *HindIII*, that excises the hygromycin resistance cassette, thereby generating the construct pM6194. A soluble construct of *msmeg_6194*, lacking 19 N-terminal and 7 C-terminal amino acids, was amplified from pD6194 and sub-cloned into the pET-28a vector at *Nde*I and *Hin*dIII sites, generating pETs6194 for protein expression and purification. Plasmids pD6194, pM6194, and pETs6194, which contain the cloned gene, were mutated from valine 139 to glutamic acid using QuikChange site-directed mutagenesis kit (Agilent Technologies, CA, USA), resulting in pD6194-V139E, pM6194-V139E, and pETs6194-V139E, respectively. All primers used are listed in Table S2 (supplemental material). All plasmid constructs and mutations were sequenced for confirmation (Eurofins Scientific, Bangalore, India).

### Deletion of the msmeg_6194 gene from M. smegmatis

The *msmeg_6194* gene was deleted from *M. smegmatis* using the homologous-recombination method (Pandey et al. 2018). The upstream and downstream flanking regions (500 bp each) of *msmeg_6194* were amplified from the *M. smegmatis* mc^2^155 chromosomal DNA (see Table S2 for primer sequences). The amplicon was cloned sequentially into the mycobacterial suicide vector pSMT100, on either side of the hygromycin cassette, to produce pSD6194 (Fig. S1). It was then electroporated into an electrocompetent *M. smegmatis* mc^2^155 cell. Transformants were selected on 7H11 plates with hygromycin and 10% (wt/vol) sucrose. Only double-crossover mutants could grow on sucrose hygromycin plates due to the sacB gene cassette in pSMT100. The deletion was confirmed by amplifying the DNA product with various primers (Fig. S1).

### Microscopic examination of the ectopic expression of MSMEG_6194 and its mutant(V139E) in a septuple PBP mutant of E. coli

The *E. coli* septuple PBP mutant CS703-1 (Ghosh & Young 2003) was transformed with the plasmids pD6194 and pD6194-V139E and cultured overnight in LB medium containing chloramphenicol (20 mg/mL) at 37°C. The overnight culture was diluted to 1% and further grown to an optical density (OD_600nm_) of 0.2, followed by induction with 0.02% (w/v) arabinose for 2 hours. Samples (5µL) were deposited on 1% agarose pad (5mm x 5mm) and visualised using phase contrast microscopy at a magnification of 100x with an Olympus iX73 microscope. For controls, CS703-1 cells containing pBAD18-Cam were used as a negative control, while pPJ5, which expresses *E. coli* PBP5, served as the positive control and was induced with 0.0005% arabinose.

### Determination of Minimum Inhibitory Concentration (MICs)

The minimum inhibitory concentration (MIC) of various antibiotics was determined following CLSI guidelines. For *E. coli,* experiments were conducted in microtitre plates with an assay volume of 300 µL per well and an inoculum size of 10^5^ cells*. E. coli* strains SK2O56-3 and AM1OC-1, transformed with plasmids pD6194 and pD6194-V139E, served as experimental samples, while a cell containing plasmid pBAD18-Cam was used as the control. Gene expression was induced by adding 0.02% (w/v) arabinose to the cultures in Mueller-Hinton broth. For *M. smegmatis*, the final well volume was adjusted to 200 µL using 7H9 medium, and gene expression was induced with 20 ng/mL tetracycline. Approximately 10^5^ cells were inoculated per well. Plates were incubated at 37 °C for 12–16 h (*E. coli*) or 48–72 h (*M. smegmatis*). Bacterial growth was measured at 600 nm using a Multiskan Spectrum spectrophotometer. Each experiment was repeated six times, with the most reproducible results were reported.

### Protein expression and purification

The soluble (s) MSMEG_6194 and its mutant (V139E) protein were overexpressed using the pETs6194 and pETs6194-V139E vectors in *E. coli* BL21 (DE3), respectively. The protein expression was induced at an optical density (OD_600nm_) of 0.4 by adding 0.5 mM IPTG to cultures grown at 37 °C, followed by incubation for 16 hours at 20 °C. The bacterial culture was harvested, lysed through sonication, and centrifuged to isolate the soluble fraction in the supernatant. The cytosolic proteins were purified via affinity chromatography (Ni²⁺-NTA) using an Akta-Prime Plus instrument (Wipro GE Healthcare, NJ, USA) and eluted with a 50–500 mM imidazole gradient in 10 mM Tris-HCl buffer (pH 7.9) containing 300 mM NaCl. The purity of the protein was verified through 12% sodium dodecyl sulphate-polyacrylamide gel electrophoresis (SDS-PAGE) (Fig. 1A). The purified proteins were then subjected to dialysis for 12 hours at 4°C, with three buffer exchanges performed without the addition of imidazole. Subsequently, it was stored at −80°C in buffer solution containing 50% glycerol for future applications.

**Figure 1.**
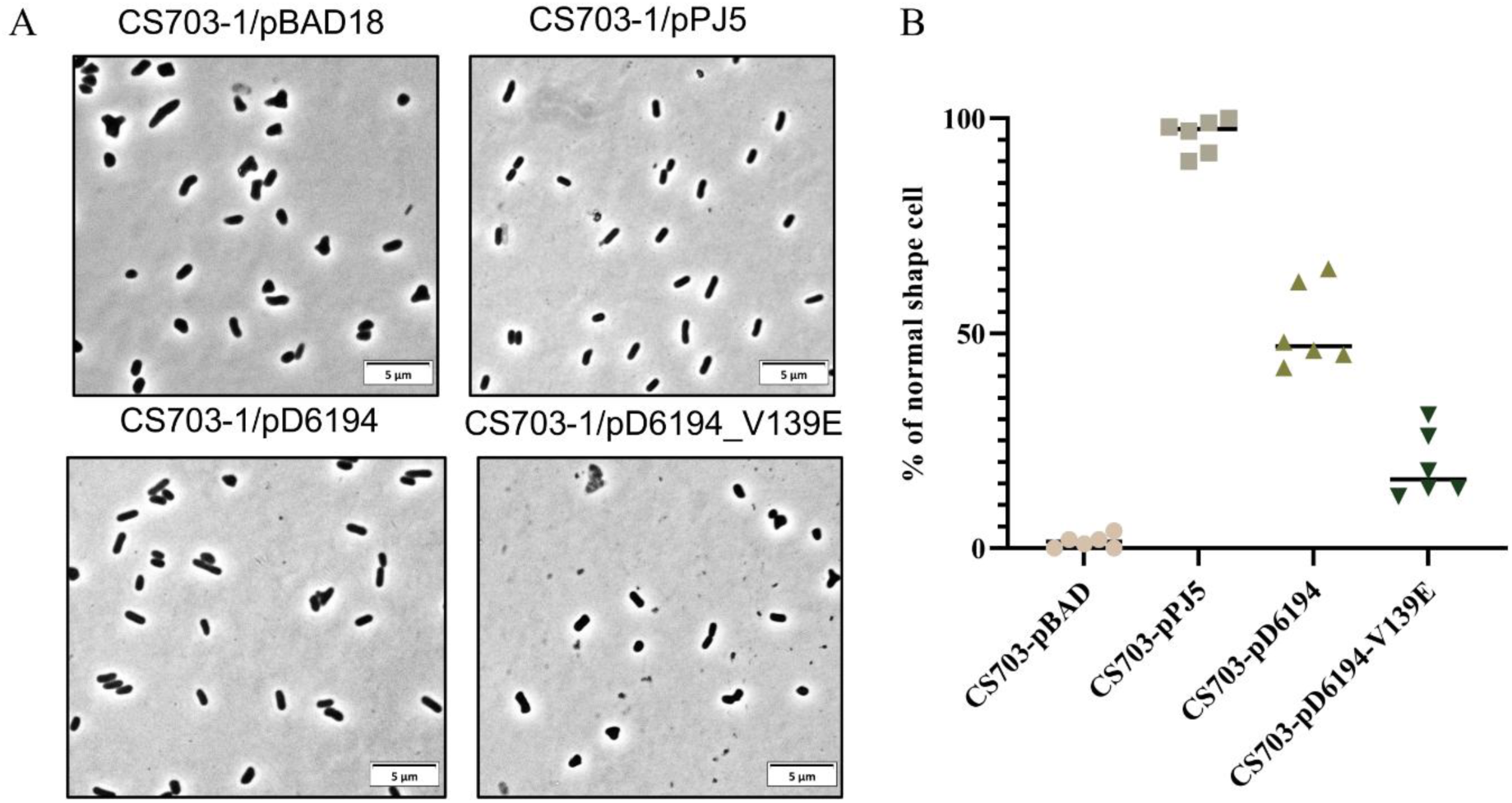
Restoration of normal cell morphology in *E. coli* CS703-1 by MSMEG_6194 and its V139E mutant. (A) The proteins PBP5 (pPJ5; positive control), MSMEG_6194 (pD6194), and the V139E mutant (pD6194-V139E) were ectopically expressed in *E. coli* CS703-1 using the pBAD18-Cm vector under arabinose induction. Cell morphology was analysed during the early logarithmic growth phase. (B) The proportion of morphologically normal cells was quantified from randomly selected microscopic fields. Each data point represents an independent experiment, and the horizontal bar indicates the mean value.

### Bocillin-FL binding assay

The functionality of the purified proteins was evaluated by labelling them with the fluorescent penicillin, Bocillin-FL (Invitrogen), in a 10 mM phosphate buffer at pH 7.4. The reaction was conducted at 35°C for 15 minutes. After denaturation by boiling, the proteins were analysed using 12% SDS-PAGE and visualised with a Typhoon FLA 7000 (GE Healthcare) at an excitation wavelength of 488 nm and an emission wavelength of 526 nm.

### Determination of acylation rate constant

The acylation rate constant was determined by incubating 25 μg of the enzyme with varying concentrations of Bocillin-FL (25, 50, and 100 μM) in a 10 mM phosphate buffer at pH 7.4. Samples were drawn at different intervals, ranging from 5 minutes to 30 minutes. A sample buffer containing SDS and beta-mercaptoethanol was added to stop the reaction. The samples were boiled and analysed using 12% SDS-PAGE and visualised with a Typhoon FLA 7000 (GE Healthcare). The intensity of the Bocillin-FL-bound protein bands was quantified through densitometric scanning using ImageJ software (Schneider *et al*., 2012). The second-order rate constant (*k_2_/K*) was calculated by determining the pseudo-first-order rate constant (*k_a_*) using the equation:

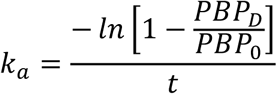

’PBP_D_’ represents the intensity of Bocillin-FL bound to protein at time *t* for a given Bocillin-FL concentration, and ‘PBP_0_’ denotes the density at enzyme saturation with Bocillin-FL. These *k_a_* values were plotted against their respective Bocillin-FL concentrations to determine *k_2_/K*.

### Determination of the deacylation rate constant

The deacylation rate constant was determined by incubating 50 μg of protein with Bocillin FL (50 μM) for 20 minutes at 35 °C in a 10 mM phosphate buffer at pH 7.4. Following this incubation, an excess of Penicillin G (3mM) was added to the mixture. The quantity of Bocillin FL remaining bound to the protein was evaluated by taking aliquots at various intervals (0-30 min). The intensity of the labelled protein was assessed using densitometric scanning after separation by 12% SDS-PAGE. The deacylation rate constant (*k_+3_*) was calculated by measuring the fluorescence of the remaining protein-bound Bocillin FL (the acyl-enzyme complex), which diminished over time.

### Determination of Steady-state kinetic parameters

The beta-lactamase activity of the purified protein was determined by measuring the changes in substrate absorbance using a UV-VIS spectrometer (Eppendorf, Germany). The wave length (λ) and the extension coefficient (ε) used for different antibiotics are given in the supplementary table S3. The hydrolysis reactions were conducted at 25 °C in 10 mM Tris-CL buffers at pH 7.4, supplemented with 150 mM NaCl, using enzyme (0.37 μM) and various beta-lactam substrates (20–500 mM). Initial reaction rates were determined from the linear portion of the hydrolysis curve. Michaelis–Menten constants (*K_m_* and *V_max_*) were obtained by fitting initial rate versus substrate data to the Michaelis–Menten equation using an in-house Python script. The turnover number (*k_cat_*) and catalytic efficiency (*k_cat_/K_m_*) of the enzyme were also calculated. Each hydrolysis reaction was repeated in triplicate for accuracy.

### In-silico analysis

The structural model of MSMEG_6194 was obtained from the AlphaFold database (Varadi et al. 2022), and the V139E mutant was generated using the MODELLER 10 software (Webb & Sali 2016). Molecular docking of the pentapeptide substrate (L-Ala–D-Glu–mDAP–D-Ala–D-Ala) was carried out with the ENZYME-DOCKER module of the CHARMM-GUI server using energy-minimised protein structures (Lee et al. 2016). Apo proteins and protein–peptide complexes were then prepared through CHARMM-GUI and subjected to 500 ns molecular dynamics simulations in GROMACS 2024.2. A detailed computational workflow is provided in the Supplementary Methods.

## Results

### MSMEG_6194 is a penicillin-interacting enzyme that shows structural similarity to class A beta-lactamase

MSMEG_6194 is annotated as a putative beta-lactamase in the UniProt database. AlphaFold3 structural modelling (Fig. S2A) and structural comparisons using the FoldSeek server (van Kempen et al. 2024) indicate similarity to TEM-type beta-lactamases, and BLASTp search against the beta-lactamase database (Naas et al. 2017) suggests similarity to the class A beta-lactamase PSE-4. However, MSMEG_6194 lacks the conserved glutamic acid in the omega-loop, a key residue for beta-lactamase function. Interestingly, phylogenetic analysis shows that MSMEG_6194 is more closely related to *E. coli* PBP5 than to TEM-type beta-lactamases (Fig. S2B).

To evaluate the functional significance of the missing catalytic residue, a glutamic acid was introduced at position 139 (V139E), corresponding to the catalytic Glu found in the omega-loop of TEM-1 beta-lactamase. The absence of this residue in MSMEG_6194 suggests that the protein is unlikely to function as a beta-lactamase in its native form. In parallel, both the native and mutant proteins were assessed for DD-carboxypeptidase activity, owing to their phylogenetic relatedness to *E. coli* PBP5.

### MSMEG_6194 partially restore the morphological defect in an E. coli septuple PBP mutant (CS703-1)

To investigate the DD-Carboxypeptidase activity of MSMEG_6194 and its mutant (V139E) in vivo, we assessed the morphological changes in CS703-1 cells following the ectopic expression of both the native and mutant proteins. *E. coli* 703-1 is derived from *E. coli* CS109), which lacks seven non-essential penicillin-binding proteins (PBPs) and displays morphological deformities (Nelson & Young, 2001, Ghosh *et al*., 2006). The trans-expression of PBP5 and other DD-carboxypeptidases in this deletion mutant restores the cell shape to near-normal, rod-like forms (Chowdhury *et al*., 2012). PBP5 from *E. coli* served as a positive control. Expression of MSMEG_6194 partially restored a uniform morphological phenotype in the aberrantly-shaped *E. coli* CS703-1, with approximately 45% to 48% of cells reverted to normal shape (Fig. 1A). In contrast, cells expressing the MSMEG_6194_V139E mutated protein showed a significant impairment in restoring their normal shape, resulting in only 18% to 22% of cells returning to their normal shape (Fig. 1B). These findings suggest that MSMEG_6194 may exhibit relatively weak DD-Carboxypeptidase activity, which is substantially hindered due to V139E mutation.

### MSMEG_6194 possesses in vitro DD-Carboxypeptidase activity

Since the cells expressing *msmeg_6194* showed considerable ability to revert the morphological oddities of CS703 cells, it was imperative to determine the DD-Carboxypeptidase activity of MSMEG_6194. To assess whether MSMEG_6194 and its V139E mutant exhibit DD-Carboxypeptidase activity *in vitro*, we investigated the ability of the respective purified proteins to release free D-alanine from two substrates: the synthetic tripeptide Nα, Nε-diacetyl-L-lysine-D-alanine-D-alanine (KAA) and the peptidoglycan-mimetic pentapeptide L-alanine-γ-D-glutamic acid-L-lysine-D-alanine-D-alanine (AEKAA). Both the wild-type enzyme and the mutant variant were capable of hydrolysing the tri- and pentapeptide substrates (Table 1), with the pentapeptide demonstrating greater efficiency for both proteins. However, the V139E mutation resulted in a notable decrease in catalytic efficiency (*k_cat_/K_m_*)—approximately 4.5-fold for the tripeptide and 12-fold for the pentapeptide. This decline was primarily attributed to a significant increase in *K_m_*, which indicates reduced substrate affinity caused by the V139E mutation. These results confirm that MSMEG_6194 possesses DD-Carboxypeptidase activity, though the activity is significantly diminished by the introduction of the V139E mutation, thus aligning with our *in vivo* observation of the reduced restoration of cell shape.

**Table 1.**
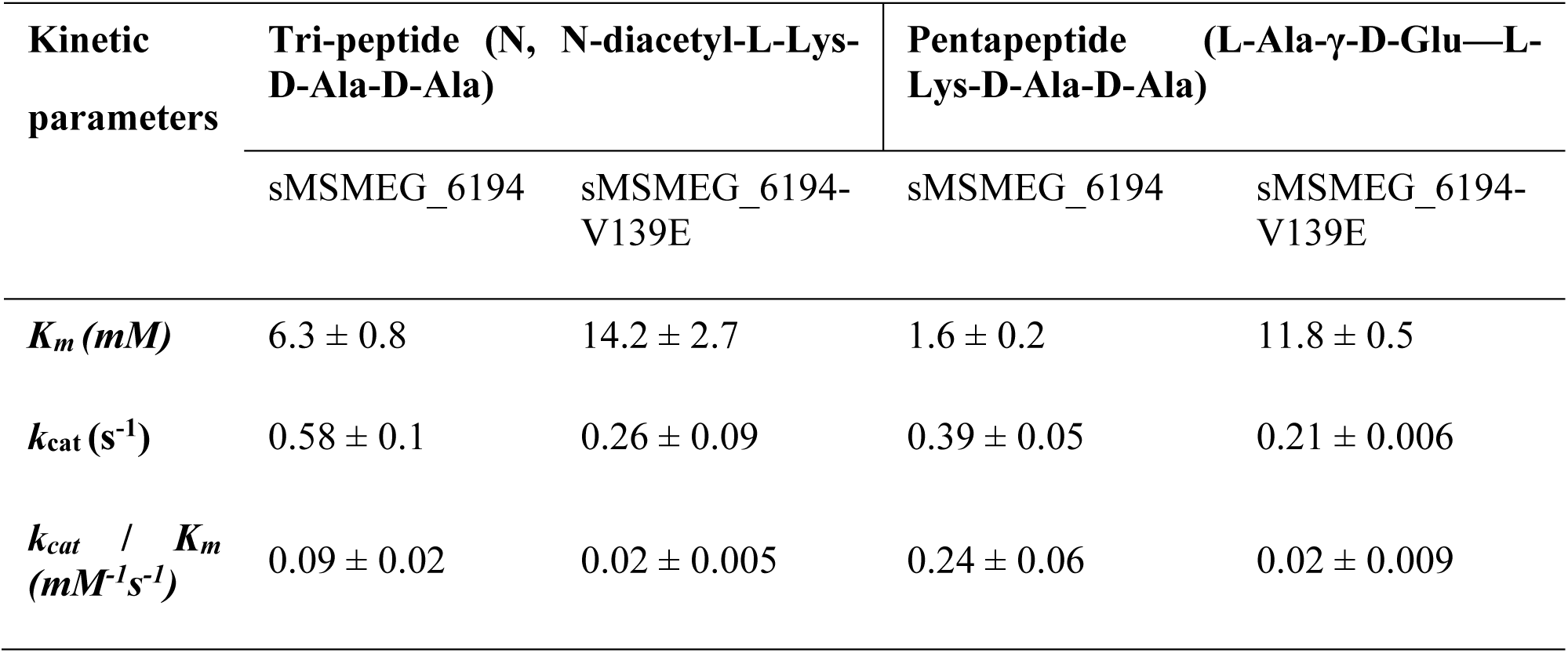
Kinetic parameters of DD-CPase activity of the sMSMEG_6194 and its mutant (V139E) proteins with peptide substrates.

| Kinetic parameters | Tri-peptide (N, N-diacetyl-L-Lys-D-Ala-D-Ala) | | Pentapeptide (L-Ala-γ-D-Glu—L-Lys-D-Ala-D-Ala) | |
| --- | --- | --- | --- | --- |
|  | sMSMEG_6194 | sMSMEG_6194-V139E | sMSMEG_6194 | sMSMEG_6194-V139E |
| $K_m$ (mM) | $6.3 \pm 0.8$ | $14.2 \pm 2.7$ | $1.6 \pm 0.2$ | $11.8 \pm 0.5$ |
| $k_{cat}$ (s<sup>-1</sup>) | $0.58 \pm 0.1$ | $0.26 \pm 0.09$ | $0.39 \pm 0.05$ | $0.21 \pm 0.006$ |
| $k_{cat} / K_m$ (mM<sup>-1</sup>s<sup>-1</sup>) | $0.09 \pm 0.02$ | $0.02 \pm 0.005$ | $0.24 \pm 0.06$ | $0.02 \pm 0.009$ |

### The V139E Substitution Improves the Deacylation Efficiency of MSMEG_6194

It was observed that both native and mutant proteins bound Bocillin-FL, confirming that MSMEG_6194 is a penicillin-interacting protein (Fig. 2A). While native MSMEG_6194 did not hydrolyse nitrocefin, the V139E variant exhibited clear beta-lactamase activity as revealed by the change in colour of nitrocefin to brown (Fig. 2B). To determine whether the V139E mutation alters the kinetic behaviour of MSMEG_6194, binding efficiency and catalytic steps were examined by characterising acyl-enzyme complex formation and measuring acylation (*k_2_/K*) and deacylation (*k ₊₃*) rate constants using Bocillin-FL. The rate constants of the wild-type and mutant proteins were quantified and compared. The acylation rate of the mutant (79.29 ± 8.3 M ⁻¹s ⁻¹) was similar to that of MSMEG_6194 (68.0 ± 9.2 M⁻¹s ⁻¹; p > 0.05), indicating no significant difference in acyl-enzyme formation. In contrast, the V139E mutant showed a markedly higher deacylation rate (33.6 ± 8.3 s ⁻¹) than the wild-type protein (12.1 ± 5.2 s ⁻¹), representing an approximately 177% increase and demonstrating faster release of the beta-lactam from the acyl-enzyme complex. This enhanced deacylation suggests a beta-lactamase-like activity for MSMEG_6194-V139E, consistent with the behaviour of beta-lactamases that rapidly cleave and release substrates. Together with the nitrocefin hydrolysis assays, these findings indicate that the V139E mutation confers beta-lactamase-like activity to MSMEG_6194, whereas the wild-type protein perhaps does not function as a beta-lactamase under the tested conditions.

**Figure 2.**
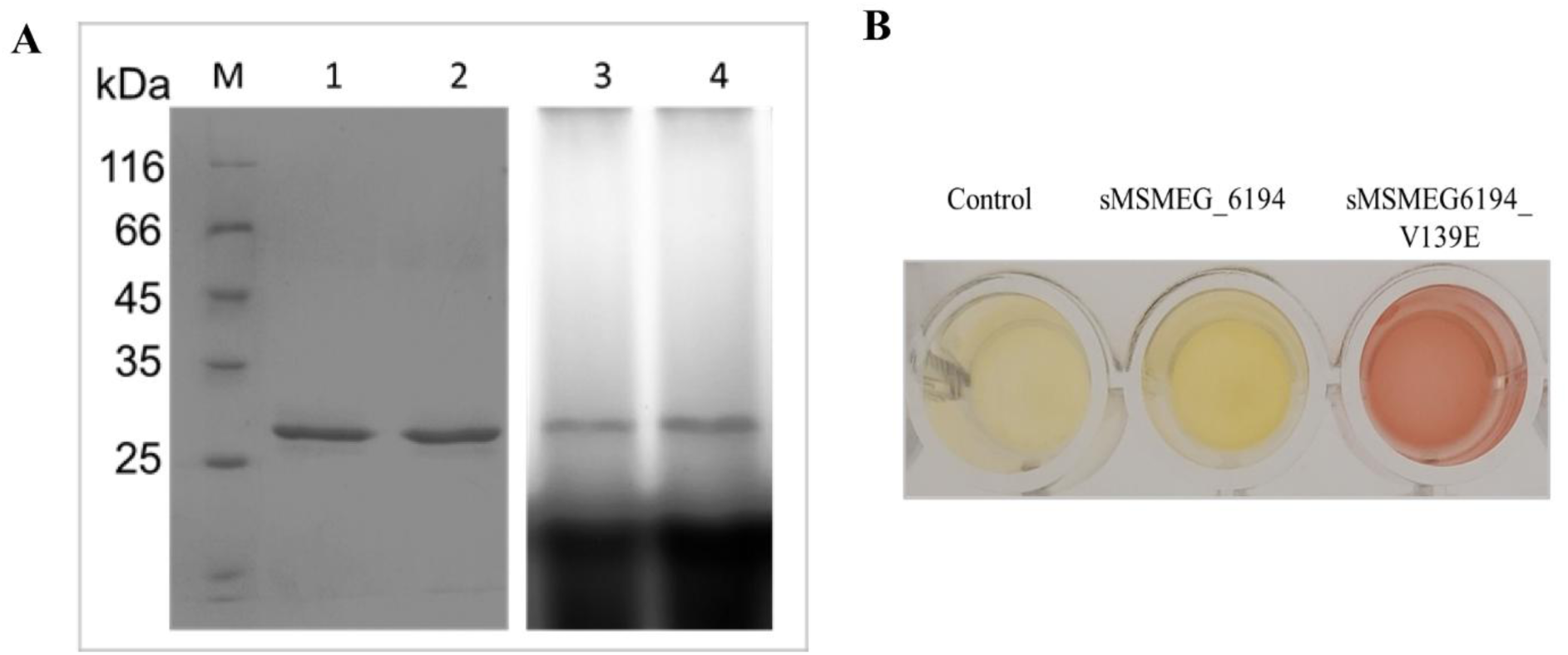
Bocillin-FL binding and Nitrocefin hydrolysis assay of sMSMEG_6194 and its mutant (V139E). (A) Fluorescent penicillin (Bocillin-FL, 25 µM) binding assay of purified sMSMEG_6194 and its V139E mutant (5 µg each). Lane 1: purified sMSMEG_6194; Lane 2: purified sMSMEG_6194-V139E; Lane 3: sMSMEG_6194 labelled with Bocillin-FL; Lane 4: sMSMEG_6194-V139E labelled with Bocillin-FL. (B) Nitrocefin hydrolysis assay of sMSMEG_6194 and its V139E mutant (1 µg each) using 100 µM nitrocefin as substrate.

### V139E substitution in MSMEG_6194 increases beta-lactam tolerance in both E. coli and M. smegmatis

To assess whether MSMEG_6194 and its mutant variant (V139E) modulate beta-lactam susceptibility, minimum inhibitory concentrations (MICs) were measured in two *E. coli* surrogate strains with increased sensitivity to beta-lactams (Table 2). *E. coli* AM10C-1 (Δ*ampC*), which lacks the beta-lactamase AmpC, and *E. coli* SK2056-3 (Δ*pbp5 Δpbp6*), which is devoid of the DD-carboxypeptidases PBP5 and PBP6, were selected to uncover potential beta-lactamase- or DD-Carboxypeptidase–like functions of MSMEG_6194, as suggested by previous studies (Pandey *et al*., 2020, Panda *et al*., 2024).

**Table 2.** Beta-lactam susceptibility (MIC, µg/mL) of *E. coli* and *M. smegmatis* strains was assessed following ectopic expression of MSMEG_6194 or its mutant V139E. Values in parentheses denote the fold change in MIC relative to strains containing an empty vector.

| **Antibiotics** | ***E. coli***<br>**2443** | ***E. coli***<br>**SK2056-1** | ***E. coli***<br>**SK2056-1/**<br>**pD6194** | ***E. coli***<br>**SK2056-1/**<br>**pD6194-**<br>**V139E** | ***E. coli***<br>**AM1OC-1** | ***E. coli***<br>**AM1OC-1/**<br>**pD6194** | ***E. coli***<br>**AM1OC-1/**<br>**pD6194-**<br>**V139E** |
| --- | --- | --- | --- | --- | --- | --- | --- |
| Ampicillin | 16 | 4 | 8 | 16 | 2 | 4 | 8 |
| Amoxicillin | 4 | 2 | 2 | 4 | 2 | 2 | 4 |
| Piperacillin | 2 | 0.5 | 0.5 | 0.5 | 0.5 | 1 | 2 |
| Ticarcillin | 8 | 0.5 | 2 | 4 | 1 | 2 | 8 |
| Cephalothin | 8 | 8 | 8 | 16 | 2 | 2 | 8 |
| Cefaclor | 8 | 4 | 4 | 4 | 0.5 | 0.5 | 2 |
| Cefoxitin | 2 | 0.5 | 0.5 | 0.5 | 0.25 | 0.25 | 0.5 |
| **Antibiotics** | ***M. smegmatis*** | ***M. smegmatis***<br>**(Δmsmeg_6194)** | ***M. smegmatis***<br>**(Δmsmeg_6194) +**<br>**pM6194** |  | ***M. smegmatis***<br>**(Δmsmeg_6194) +**<br>**pM6194-V139E** |  |  |
| Ampicillin | 512 | 512 | 512 |  | 1024 |  |  |
| Ticarcillin | 64 | 64 | 64 |  | 256 |  |  |
| Cefoxitin | 64 | 64 | 64 |  | 128 |  |  |

Ectopic expression of MSMEG_6194 in *E. coli* SK2056-3 (Fig. S3) resulted in a moderate increase in resistance for several penicillin-groups of antibiotics, including a 2-fold increase for ampicillin and a 4-fold increase for ticarcillin. This moderate elevation indicates that MSMEG_6194 can provide protection against these cell wall-targeting antibiotics. Importantly, the V139E mutant conferred a more pronounced resistant phenotype, displaying higher fold changes relative to the vector control, including 4-fold for ampicillin, 2-fold for amoxicillin, 8-fold for ticarcillin, and 2-fold for cephalothin. These observations indicate that the V139E substitution enhances the beta-lactam protective function of MSMEG_6194.

In the beta-lactamase–deficient *ΔampC* strain, MSMEG_6194 alone caused only a mild 2-fold increase in MICs for ampicillin, piperacillin, and ticarcillin. In contrast, the V139E variant showed a marked increase in resistance across multiple antibiotic classes, with 4-, 2-, 4-, and 8-fold increases observed for ampicillin, amoxicillin, piperacillin, and ticarcillin, respectively, and 4-, 4-, and 2-fold increases for cephalothins, cefaclor, and cefoxitin, respectively. Although the native protein displayed only limited activity, the substantially higher MICs conferred by V139E mutant, together with its increased deacylation efficiency, strongly support a gain-of beta-lactamase-like function in MSMEG_6194.

To validate these findings in the native host, beta-lactam resistance profiles were assessed in *M. smegmatis* (Table 2). Deletion of *msmeg_6194* did not alter susceptibility compared to the wild-type strain, and ectopic expression of the native gene did not change MICs either. However, expression of the V139E mutant increased resistance, as evidenced by 2-fold increases for ampicillin, 4-fold increases for ticarcillin, and 2-fold increases for cefoxitin relative to the cell harbouring the vector control.

Collectively, these results demonstrate that MSMEG_6194 has a modest protective role against beta-lactams, possibly due to antibiotic masking (Sarkar et al. 2008) instead of antibiotic hydrolysis, while the V139E mutation significantly enhances the hydrolysing activity. The stronger resistance phenotype observed in beta–lactamase–deficient *E. coli* and in *M. smegmatis* expressing V139E suggests that this mutation may confer beta-lactamase-like function, thereby enhancing survival mycobacteria against diverse beta-lactam antibiotics.

### The V139E mutation confers penicillinase-like beta-lactamase activity to sMSMEG_6194

To determine the functional impact of the V139E substitution, steady-state kinetic assays were performed using purified sMSMEG_6194-V139E against a panel of six beta-lactam antibiotics, including penicillins and first-generation cephalosporins. Under identical conditions, the wild-type MSMEG_6194 exhibited no detectable hydrolytic activity toward any substrate, confirming that the V139E substitution was essential for the acquisition of beta-lactamase activity. The kinetic parameters derived from Michaelis–Menten analysis are summarised in Table 3.

**Table 3.** Kinetic parameter for the beta-lactam activity of purified sMSMEG_6194 -V139E mutant protein. The data are presented as the mean ± SD from three independent replicates.

| Antibiotics | $K_m$ ($\mu M$) | $k_{cat}$ ($s^{-1}$) | $k_{cat} / K_m$ ($\mu M^{-1} s^{-1}$) $\times 10^{-3}$ |
| --- | --- | --- | --- |
| Ampicillin | 186.6 ± 25.8 | 4.86 ± 0.18 | 26.0 ± 3.0 |
| Amoxicillin | 326.3 ± 49.6 | 0.81 ± 0.02 | 2.5 ± 0.3 |
| Piperacillin | 234.7 ± 21.36 | 3.5 ± 0.12 | 14.9 ± 9 |
| Ticarcillin | 163.13 ± 30.17 | 9.2 ± 1.01 | 56.4 ± 10.2 |
| Cephalothin | 289.66 ± 71.21 | 0.78 ± 0.16 | 2.7 ± 0.1 |
| Cefaclor | 335.93 ± 57.7 | 0.49 ± 0.06 | 1.4 ± 0.3 |

Among the tested substrates, ticarcillin showed the highest catalytic efficiency (*k_cat_/K_m_*), followed by ampicillin and piperacillin. This elevated efficiency resulted from both the high turnover rate (*k_cat_*) and strong substrate affinity (low *K_m_*). In contrast, cephalosporins (cephalothin and cefaclor) and amoxicillin displayed only moderate efficiencies, primarily due to reduced turnover rates. With the exception of amoxicillin, all cephalosporins demonstrated substantially lower catalytic efficiency compared with the penicillin class. Notably, cephalothin exhibited an approximately 19-fold decrease in activity relative to ticarcillin. Overall, these results indicate that the V139E substitution confers beta-lactamase activity on sMSMEG_6194 with a clear substrate preference for penicillin-class antibiotics—particularly carboxy-penicillins such as ticarcillin, while activity toward cephalosporins remains limited. This pattern suggests that the V139E mutant functions similarly to a carboxy-penicillinase-type beta-lactamase.

### MD simulation reveals that the V139E mutation destabilises the protein–pentapeptide interaction

Our enzymatic assays confirmed that MSMEG_6194 exhibits DD-carboxypeptidase activity, which was markedly reduced upon introduction of the V139E mutation. To elucidate the structural basis of this loss of activity, molecular dynamics (MD) simulations were performed for both the native and mutant proteins in their apo forms and in the docked complex with a pentapeptide substrate (Fig. S4). Both apo-proteins reached structural stability after approximately 100 ns, whereas the protein–peptide complexes stabilised more slowly—the wild-type after ∼200 ns and the mutant after ∼380 ns—indicating delayed equilibration of the mutant complex. RMSF and principal component analyses revealed enhanced flexibility in residues 198–209 and within the omega-loop region (residues 136–151), the latter being proximal to the mutation site, suggesting local destabilisation induced by V139E (Fig. S5).

A particularly striking difference was observed in substrate behaviour within the active site (Fig. 3). In the wild-type complex, the pentapeptide substrate remained stably bound, maintaining consistent interactions between its terminal D-Ala-D-Ala moiety and the catalytic residues throughout the simulation. In contrast, in the V139E mutant, the substrate began to dissociate after ∼120 ns, indicating weakened substrate retention. As both systems were modelled under identical conditions, this divergence reflects a genuine mutation-driven effect rather than artefacts of modelling. Together, these findings provide a structural rationale for the experimentally observed loss of DD-Carboxypeptidase activity in the V139E mutant of MSMEG_6194, linking local conformational destabilisation to impaired substrate binding and catalysis.

**Figure 3.**
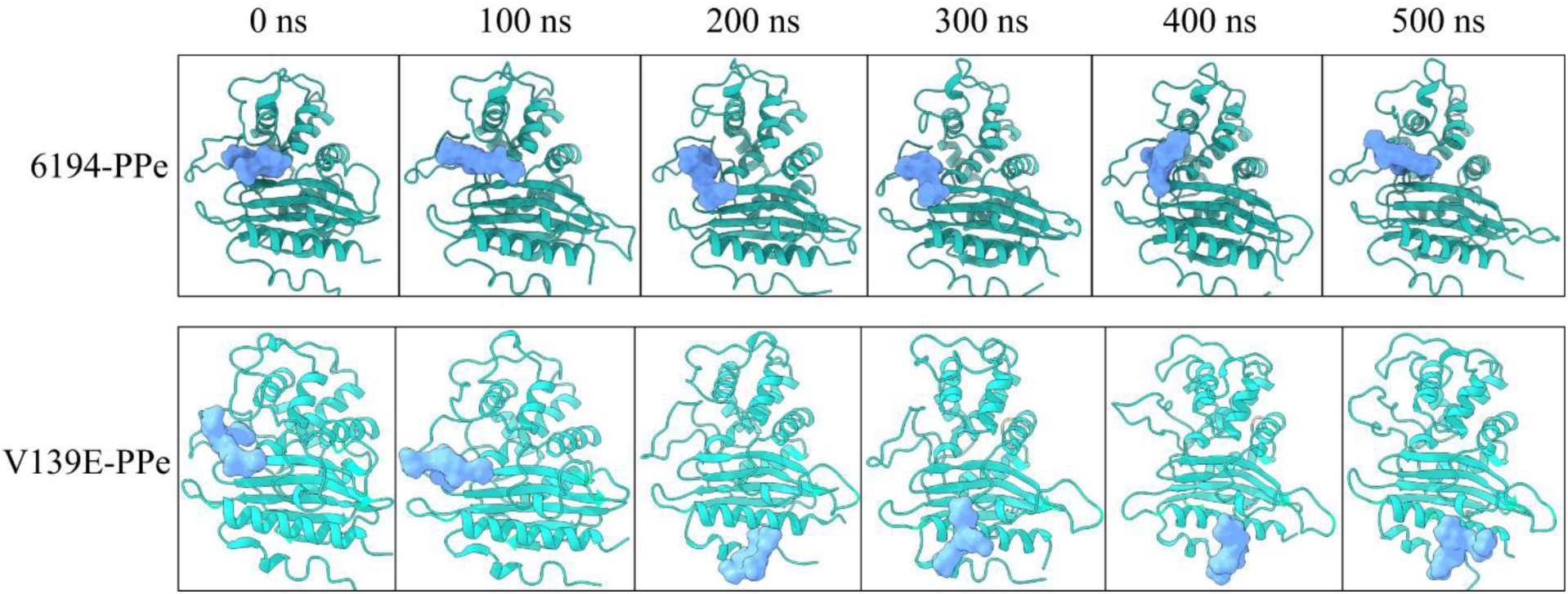
Structural dynamics of the peptide substrate within the active-site binding pocket of MSMEG_6194 and its V139E mutant during molecular dynamics simulations. Representative snapshots at different simulation intervals (0–500 ns) are shown for the wild-type MSMEG_6194–pentapeptide complex (top panel) and the V139E mutant complex (bottom panel). The protein backbone is depicted as a cyan cartoon, and the peptide substrate is shown as a blue surface, illustrating the conformational changes and substrate stability over the simulation period.

## Discussions

*Mycobacterium smegmatis* harbours a broader repertoire of penicillin-interacting enzymes. The coexistence of dual-function enzymes with both beta-lactamase and DD-carboxypeptidase activities, along with extended-spectrum class C beta-lactamases, reflects the evolutionary diversification of beta-lactamases and the emergence of adaptive beta-lactam resistance mechanisms in mycobacteria (Bansal et al. 2015, Pandey et al. 2020, Panda et al. 2025). In this study, we functionally characterised MSMEG_6194, a gene annotated as a beta-lactamase-related protein and structurally similar to class A beta-lactamases. Interestingly, despite this structural resemblance, MSMEG_6194 exhibited no detectable beta-lactamase activity under the tested conditions, though it demonstrated clear, though moderate DD-carboxypeptidase activity both *in vitro* and *in vivo*.

This observation indicates a possible evolutionary retention of DD-Carboxypeptidase function despite the acquisition of beta-lactamase-like structural features. The evolutionary transition between penicillin-binding proteins (PBPs) and beta-lactamases has been linked to alterations in two loop regions near the active site: one governing dipeptide orientation and the other facilitating the hydrolysis of the terminal peptide bond. In beta-lactamases such as TEM-1, shortening of the first loop and extension of the second (forming the “omega-loop”) promote beta-lactamase activity (Massova & Mobashery 1998, Egorov et al. 2019). However, MSMEG_6194 retains DD-Carboxypeptidase activity despite a TEM-1–like loop configuration, suggesting that loop length modification alone is insufficient to eliminate DD-Carboxypeptidase function. Consistent with this, enzyme kinetics reveal that MSMEG_6194 retains approximately 23% of the DD-Carboxypeptidase activity of PBP5 against an artificial penta-peptide substrate (Chowdhury et al. 2012). Comparable results have been reported for chimeric beta-lactamase-PBP enzymes that regain limited DD-Carboxypeptidase activity following the substitution of 28 amino acids from TEM-1 beta-lactamase with the corresponding residue of *E. coli* PBP-5 (Chang et al. 1990). Molecular dynamics (MD) simulations provided further mechanistic insight into the function of MSMEG_6194. The simulations revealed a stable pentapeptide-binding pocket in the native enzyme, with an arrangement similar to that observed in PBP5 (Nicholas et al. 2003). This structural conservation likely explains the underlying DD-Carboxypeptidase activity of MSMEG_6194.

In contrast, the introduction of a glutamic acid residue (V139E) at the position corresponding to Glu166 of class A beta-lactamases endowed MSMEG_6194 with measurable beta-lactamase activity. A similar observation was reported for *E. coli* PBP5 (Kar *et al*., 2018). These findings support the hypothesis that the absence of a catalytic glutamic acid within the omega-loop accounts for the lack of β-lactamase activity in the native protein. However, this gain in beta-lactamase function came at the expense of DD-Carboxypeptidase activity, suggesting a trade-off between these catalytic modes. MD simulations of the MSMEG_6194-V139E mutant revealed instability in the pentapeptide–protein complex compared to the wild-type enzyme. The introduced glutamate residue appeared to generate electrostatic repulsion with the terminal carboxyl group of the peptide substrate, impairing proper binding. This observation aligns with earlier reports, in which the introduction of glutamic acid in the omega-loop of *E. coli* PBP5 similarly reduced DD-Carboxypeptidase activity (Dutta et al. 2015).

Despite its structural similarity to TEM-1, the mutant enzyme exhibited markedly reduced beta-lactamase efficiency—approximately 10^3^ fold lower catalytic efficiency toward ampicillin than TEM-1 (Brown et al. 2009). This reduced enzymatic activity is likely due to the absence of a conserved asparagine residue (N170 in TEM-1), which normally stabilises the catalytic water molecule in concert with Glu166 (He et al. 2020). Additionally, increased flexibility of the omega-loop in the mutant protein may further contribute to the diminished activity. In class A beta-lactamases, the rigidity of the omega-loop—particularly at its N-terminal region—is critical for maintaining the optimal positioning of the catalytic Glu166 residue (Meneksedag et al. 2013). Therefore, enhanced omega-loop mobility could disrupt the precise geometric arrangement of the catalytic site, ultimately compromising the efficiency of beta-lactam hydrolysis.

Overall, our findings reveal that MSMEG_6194 represents an intriguing evolutionary intermediate between DD-carboxypeptidases and beta-lactamases. While its structural framework resembles that of class A beta-lactamases, the enzyme exerts DD-Carboxypeptidase activity but lacks efficient beta-lactamase function. The V139E substitution partially reintroduces beta-lactamase activity, though it destabilises DD-Carboxypeptidase function, highlighting a delicate balance between structural adaptation and catalytic efficiency. These insights reveal the evolutionary and mechanistic flexibility of penicillin-interacting enzymes in *M. smegmatis*, offering a useful basis for understanding how beta-lactamase activity varies among mycobacteria.

## Acknowledgements

APP, DC, and AR thank IIT Kharagpur for their fellowships. We appreciate the support and cooperation of the Molecular Microbiology Laboratory, Department of Bioscience and Biotechnology, IIT Kharagpur. Computational resources were provided by the Supercomputing Facility at IIT Kharagpur, established under the National Supercomputing Mission (NSM) of the Government of India, and supported by the Centre for Development of Advanced Computing (C-DAC), Pune.

## Author contributions

APP and ASG conceptualised and designed the study. APP, DC, and SB performed data acquisition, analysis, and interpretation. AR contributed to parts of the protein purification and kinetic experiments. APP and ASG wrote and edited the manuscript.

