## Supplement File for "Substitution of a valine to glutamic acid in the omega-like loop of MSMEG_6194 of *Mycobacterium smegmatis* interchanges its activity from DD-carboxypeptidase to beta-lactamase"

**Affiliation**

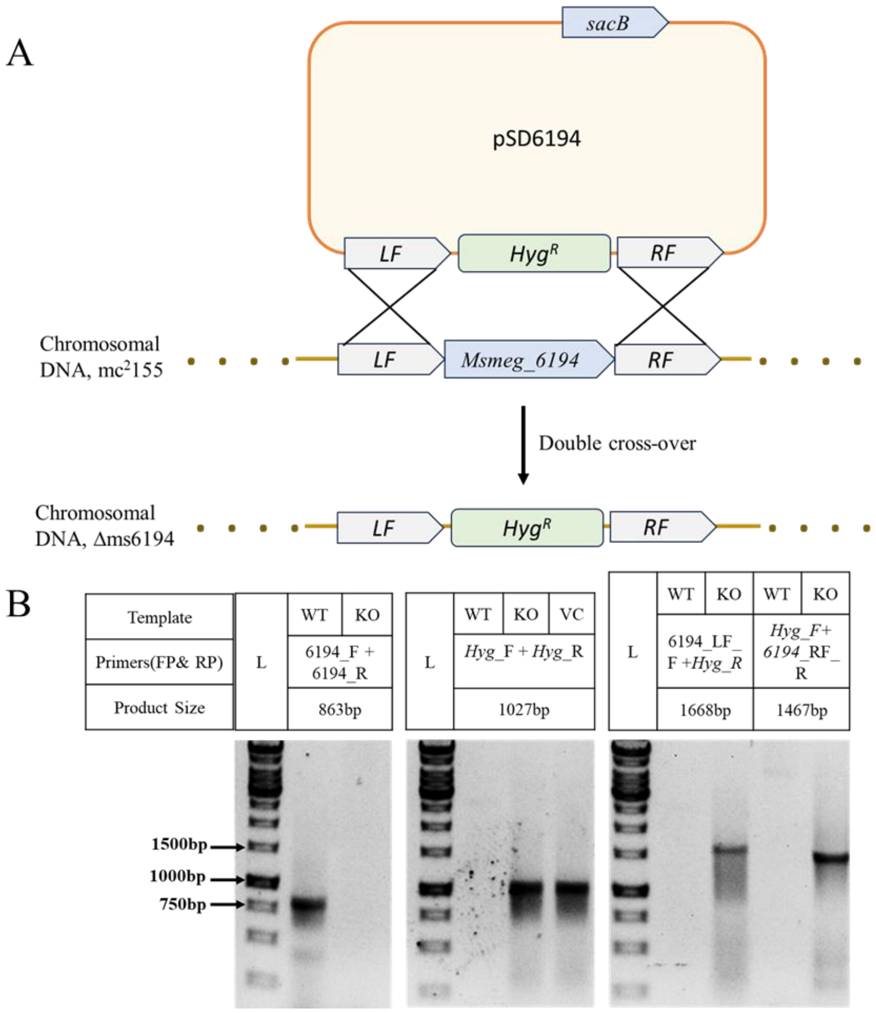


**Fig. S1. Deletion of *msmeg_6194* from the *M. smegmatis* mc²155 chromosome**. (A) Schematic representation of a double-crossover event during homologous recombination. The non-replicative plasmid pSD6194 was constructed from a mycobacterial suicide vector, pSMT100 bearing sacB and a hygromycin cassette, by stepwise cloning of the left flanking (LF) and right flanking (RF) regions of *msmeg_6194*, on both sides of the hygromycin cassette. The pSD6194 was then transformed into electrocompetent *M. smegmatis* mc²155, and the knockout mutant Δ*ms6194* was selected on a Middlebrook 7H11 plate containing hygromycin and 10% sucrose. The expression of levansucrase (from SacB) is lethal; therefore, a single crossed-over strain cannot grow on the plate. (B) Confirmation of the mutant strain by PCR using various sets of primers (refer to Table S1). The WT (mc²155), KO (Δms6194) and VC (vector control; pSMT100 harbouring hygromycin, used as positive control) were used as templates for the respective primer pairs (shown in the figure). The expected size of amplicons is shown in a 0.8% agarose gel. ‘L’ represents the DNA ladder.

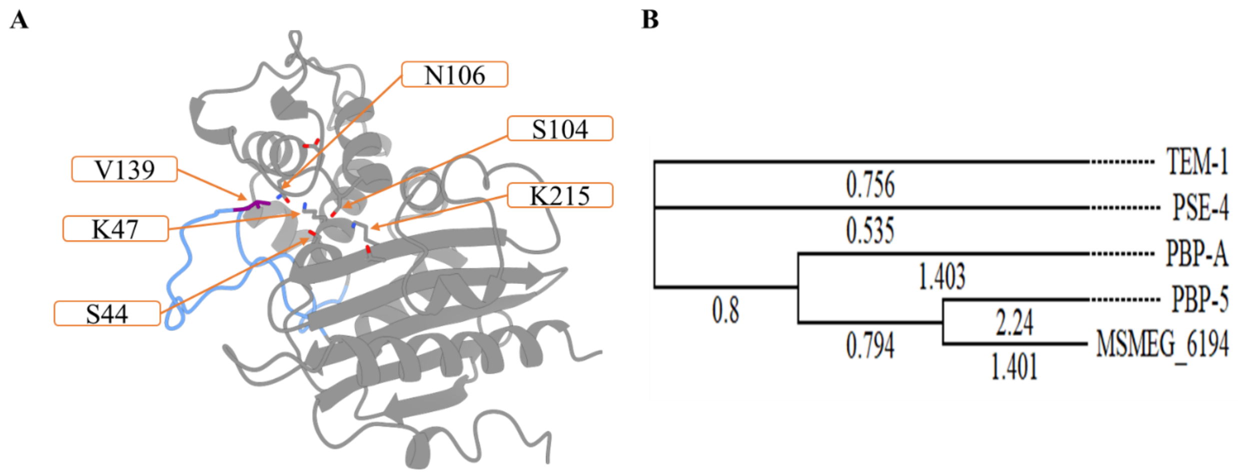


**Figure S2. Predicted structure and phylogenetic relationship of MSMEG_6194.** (A) AlphaFold-predicted structure of MSMEG_6194 (AF-A0R5H7-F1), highlighting the key active-site residues. The omega-loop region is shown in blue, with residue V139 represented as a purple side chain, which is substituted with glutamic acid in the V139E mutant. (B) Phylogenetic analysis of MSMEG_6194 and related proteins was constructed using the maximum-likelihood method. The tree includes representative beta-lactamases (TEM-1 and PSE-4) and penicillin-binding proteins PBP-A (*Thermosynechococcus elongatus*) and PBP5 (*Escherichia coli*).

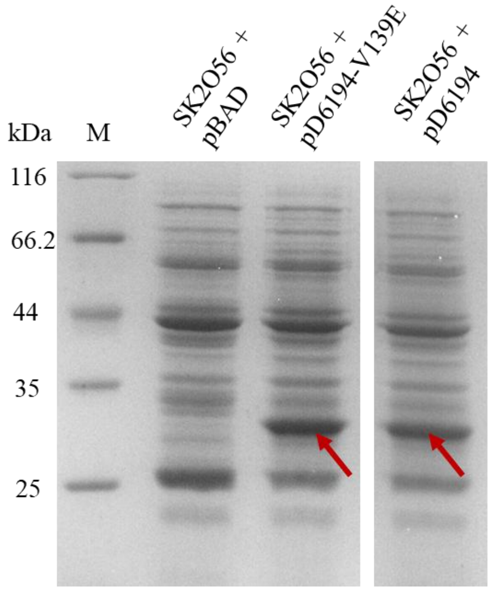


**Figure S3: Expression of MSMEG_6194 and its V139E mutant in *E. coli* SK2O56 cells.** SDS-PAGE analysis of periplasmic extracts from ~5 g of *E. coli* SK2056 cells expressing MSMEG_6194 or its V139E mutant, alongside cells carrying the empty vector (pBAD18-cm). All cultures were induced with 0.2% arabinose. Lane 'M' represents the molecular weight marker, with sizes indicated in kilodaltons (kDa). The red arrow denotes the ~30 kDa band corresponding to MSMEG_6194 and its V139E variant. (See Supporting Methods for details.)

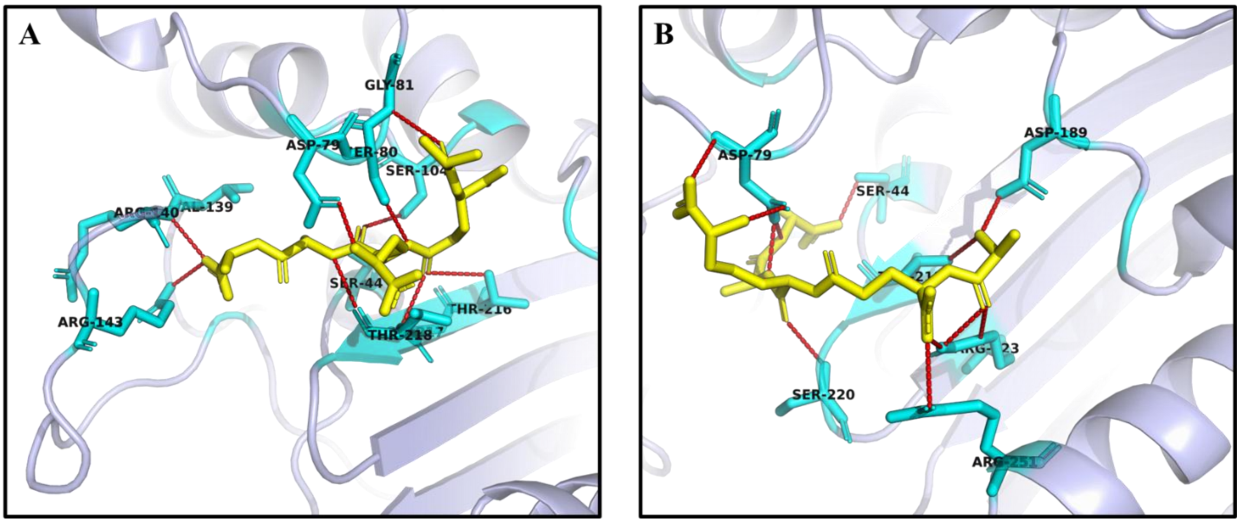


**Figure S4. Docking of the pentapeptide substrate with MSMEG_6194 and its V139E mutant.** Molecular docking of the pentapeptide substrate (L-Ala–D-Glu–mDAP–D-Ala–D-Ala) was performed with wild-type MSMEG_6194 (A) and the V139E mutant (B) using the Enzyme-Docker module of the CHARMM-GUI web server. The docked complexes were visualised in PyMOL. The peptide is shown in yellow, and hydrogen bonds are depicted as red dashed lines.

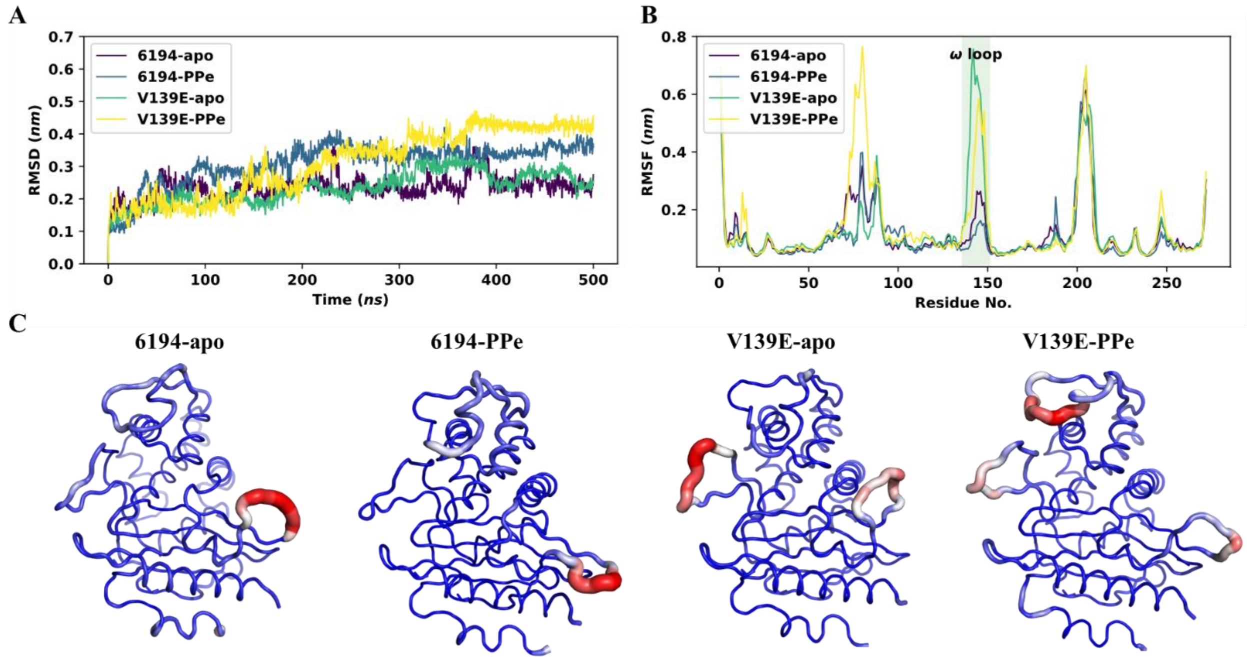


**Figure S5. Structural dynamics of MSMEG_6194 and its V139E mutant.** (A) Backbone RMSD profiles of native and mutant proteins in apo and pentapeptide (PPe)-bound states over 500 ns MD simulations. (B) RMSF plots showing residue-wise flexibility, with increased fluctuation observed in the omega-loop region of the V139E-PPe complex. (C) Residue flexibility is represented as a putty model of the first principal component (PC1), highlighting enhanced mobility of the omega-loop (red) in the mutant.

**Table S1.** Bacterial strains and plasmids used in this study.

| Strain/plasmid | Genotype/ Feature | Reference |
|---|---|---|
| **Strains** | | |
| $MC^2155$ | Wild-type strain of *M. smegmatis* | ATCC |
| *E. coli* XL1-Blue | *RecAendAhsdRsupEthirecAgyrArelAlac* | (Stratagene, LaJolla, CA) |
| *E. coli* BL21 (DE3) | $F^-$ *ompT hsdS*$_B$ ($r_B^-$ $m_B^-$) *gal dcm* (DE3) | (Stratagene, West Cedar Creek, TX) |
| *E. coli* 2443 | *thr-1 leuB6 D(gpt-proA)66 argE3 thi-1 rfb*$_{O8}$*lacY1 ara-14 galK2 xyl-5 mtl-1mgl51 rpsL31 kdgK51 supE44 Δ(recCptrrecBrecD) : : P*$_{lac}$*-bet exo(Cam*$^R$*)* | (Sandlin et al. 1996) |
| *E. coli* AM1OC-1 | 2443 Δ*ampC* | Lab stock |
| *E. coli* SK2O56-3 | 2443 Δ*pbp5pbp6* | Lab stock |
| Δ*ms6194* | *msmeg_6194* deletion strain of *M. smegmatis*; $Hyg^r$ | This study |
| *E. coli* CS703-1 | W1485 *rpoSrphΔmrcAΔdacBΔdacAΔdacCΔpbpG ΔampCΔampH* | Ghosh et al. 2003 |
| **Plasmids** | | |
| pBAD18-Cm | Expression vector with arabinose-inducible promoter; Chloramphenicol resistance ($Cm^R$) | (Guzman et al. 1995) |
| pD6194 | pBAD18-Cm, harbouring the full open reading frame (ORF) of *msmeg_6194* | This study |
| pD6194_V139E | pBAD18-Cm, harbouring the full open reading frame (ORF) of *msmeg_6194* single amino acid mutant V139E. | This study |
| pET 28a (+) | *E. coli* expression vector generating $His_6$ fusion proteins for protein over-expression; kanamycin resistance ($kan^R$) | Novagen, Madison, WI |
| pETs6194 | pET28a (+) harbouring soluble fraction of *msmeg_6194* | This study |
| pETs6194_V139E | pET28a (+) harbouring soluble fraction of *msmeg_6194* single amino acid mutant V139E | This study |
| pSMT100 | Mycobacterial suicide vector; *sacB* $Hyg^r$ | Pandey et al. 2018 |
| pSD6194 | pSMT100 with right and left flank regions of *msmeg_6194*; $Hyg^r$ | This study |
| pMIND | Tetracycline inducible vector; $Kan^r$ | Bose Institute, Kolkata, India |
| pM6194 | pMIND harbouring *msmeg_6194;* $Kan^r$ | This study |
| pM6194-V139E | pMIND harbouring *msmeg_6194* single amino acid mutant V139E*;* $Kan^r$ | This study |

**Table S2.** The following synthesised primers were used in this study.

| Primer Name | Sequence | Length | Comment |
|---|---|---|---|
| 6194_F (*NheI*) | CTCTCTGCTAGCAGGAGGCTCTCTATGAGTGACTTCTTCACCCGCG | 863bp | For cloning in pBAD18-CM |
| 6194_R (*HindIII*) | CTCTCTAAGCTTCTAGCGGTTCTGCAGGTACCGG | | |
| PM_6194_F (*NdeI*) | CTCTCTCATATGAGGAGGCTCTCTATGAGTGACTTCTTCACCCGCG | 863bp | For cloning in pMIND |
| PM_6194_F (*HindIII*) | CTCTCTAAGCTTCTAGCGGTTCTGCAGGTACCGG | | |
| s6194_F (*NdeI*) | CTCTCTCATATGGTCCGCATCACCGACCTGAACACC | 762bp | For cloning in pET-28a |
| s6194_R (*HindIII*) | CTCTCTAAGCTTCATCTCGCAGCCCAGCCATCCCAT | | |
| 6194_V139E_F | GACTGCGGGACCGGGAGCGGCTGAACCGATC | - | To perform site-directed mutagenesis to introduce a V139E point mutation. |
| 6194_V139E_R | GATCGGTTCAGCCGCTCCCGGTCCCGCAGTC | | |
| 6194_LF_F (*XmaI*) | CTCTCTCCCGGGGCTCAACTCGTCGATCTACACG | 524bp | The left flanking region of *msmeg_6194 was* cloned into the pSMT100 vector |
| 6194_LF_R (*SpeI*) | CTCTCTACTAGTCGGTTCTCGCAGCCTGTTTCG | | |
| 6194_RF_F (*PstI*) | CTCTCTCTGCAGGCGGTTTCGGTGACGCCGG | 524bp | The right flanking region of *msmeg_6194 was* cloned into the pSMT100 vector |
| 6194_RF_R (*NdeI*) | CTCTCTCATATGGGCTCCCAGATGCTCTCCGACG | | |
| *Hyg*_F | CTCTCTGGATCCATGTGTGACACAAGAATCCCTGT | 1027bp | - |
| *Hyg*_R | CTCTCTTCTAGATTAGGCGCCGGGGGCGGTGTCCG | | |

**Table S3.** Experimental parameters for kinetic studies of sMSMEG_6194-V139E protein.

| **Antibiotic** | **Δε ($M^{-1}cm^{-1}$)** | **λ (nm)** | **Antibiotic concentration (μM)** | **Enzyme concentration (μM)** |
|---|---|---|---|---|
| **Ampicillin** | -800 | **235** | **100-500** | ~0.37 |
| **Amoxicillin** | -1100 | **232** | | |
| **Piperacillin** | -1070 | **235** | | |
| **Ticarcillin** | -660 | **235** | | |
| **Cephalothin** | -6500 | **260** | **20-100** | |
| **Cefaclor** | -2670 | **264** | | |

### Supplementary Methods:

#### *Molecular Dynamics simulation:*

Molecular dynamics (MD) simulations were performed using GROMACS version 2024.2. The CHARMM36-Jul2021 force field was applied to the protein, while ligand parameters were derived from the CGenFF force field. Each system—apo or protein–ligand complex—was placed in a cubic simulation box extending at least 10 Å beyond the protein in all directions. Solvation was carried out with TIP3P water molecules, and counterions ($Na^+$ or $Cl^-$) were added to neutralise the system. Energy minimisation was performed using the steepest descent algorithm for 50,000 steps. Equilibration was carried out in two stages: a one-ns NVT simulation at 303.15 K using the v-rescale thermostat, followed by a one-ns NPT simulation at 303.15 K and 1 bar using the C-rescale barostat. During equilibration, position restraints were applied to the protein backbone and ligand heavy atoms. System stability was verified by monitoring the convergence of temperature, pressure, and density. Production MD simulations were performed for 500 ns with a 2-fs integration time step. A cutoff distance of 1.2 nm was used for short-range van der Waals interactions. Coordinates were recorded every 250 ps for subsequent analysis. Analyses were performed using tools available in GROMACS and the MDAnalysis Python package, while visualisation was carried out using PyMol. All simulations were performed on the PARAM-SHAKTI supercomputer facility available at IIT Kharagpur.

#### *Periplasmic Protein Extraction:*

Periplasmic proteins were isolated using a modified cold osmotic shock method (Malherbe et al. 2019). E. coli cultures (30 mL LB broth) were inoculated with 0.1% overnight-grown cells carrying recombinant pD6194 or its mutant (pD6194-V139E). Cultures were grown at 37°C to $OD_{600}$ ~0.2, induced with 0.2% arabinose, and incubated until late log phase. Cells were harvested (5000 × g, 5 min), washed three times with PBS (pH 7.4), and resuspended in osmotic lysis buffer (100 mM Tris-HCl, pH 8.0, 500 mM sucrose, 500 mM EDTA, 1 mg/mL lysozyme). After 15 minutes on ice, cells were centrifuged (16,000 × g, 20 min, 4°C), and the pellet was resuspended in 100 µL of distilled water. The cells were then incubated on ice for 1 minute and treated with 20 mM $Mgso_4$. Following centrifugation (16,000 × g, 20 min, 4°C), the periplasmic fraction (supernatant) was collected and analysed by SDS-PAGE.

#### *References*

Ghosh AS & Young KD Sequences near the active site in chimeric penicillin binding proteins 5 and 6 affect uniform morphology of Escherichia coli. *J Bacteriol* 2003; **185**: 2178-2186.

Guzman LM, Belin D, Carson MJ et al. Tight regulation, modulation, and high-level expression by vectors containing the arabinose PBAD promoter. J Bacteriol 1995; 177: 4121 4130

Malherbe G, Humphreys DP, Davé E. A Robust Fractionation Method for Protein Subcellular Localisation Studies in *Escherichia Coli*, *BioTechniques* 2019; **66**:171-178.

Pandey SD, Pal S, Kumar NG *et al.* Two dd-Carboxypeptidases from Mycobacterium smegmatis Affect Cell Surface Properties through Regulation of Peptidoglycan Cross-Linking and Glycopeptidolipids. *J Bacteriol* 2018; **200**.

Sandlin RC, Goldberg MB, Maurelli AT Effect of O side-chain length and composition on the virulence of Shigella flexneri 2a. Mol Microbiol 1996; 22: 63–73.
